# Space use and movement of jaguar (*Panthera onca*) in western Paraguay

**DOI:** 10.1101/119412

**Authors:** Roy T McBride, Jeffrey J Thompson

## Abstract

We estimated home range and core area size for jaguar (*Panthera onca*) in western Paraguay in the Dry Chaco, Humid Chaco and Pantanal using an autocorrelated kernel density estimator. Mean home range size was 818 km^2^ (95% CI:425-1981) in the Dry Chaco and 237 km^2^ (95% CI:90-427) in the Humid Chaco/Pantanal. Core areas, defined as the home range area where use was equal to expected use, was consistent across sexes and systems represented on average by the 59% utility distribution isopleth (range:56-64%). Males had a higher probability of larger home ranges and more directional and greater daily movements than females collectively and within systems. The large home ranges in the Dry Chaco are attributable to the relatively low productivity of that semi-arid ecosystem and high heterogeneity in resource distribution while larger than expected home ranges in the Humid Chaco/Pantanal compared to home range estimates from the Brazilian Pantanal may be due to differences in geomorphology and hydrological cycle. The large home ranges of jaguars in western Paraguay and a low proportional area of protected areas in the region demonstrate the importance of private ranchland for the long-term conservation of the species.

## Introduction

Globally, apex predators, and the maintenance of their functional roles, are severely threatened due to anthropogenic pressures, particularly associated with large spatial needs to access sufficient prey to meet metabolic requirements and persecution (Ripple et al. 2014). Habitat conversion and degradation and over hunting of prey species increase spatial requirements of apex predators, increasing conflict with humans and affecting social behavior, dispersal and habitat use (McDonald 1983; Crooks 2002; Cardillo et al. 2004; Ripple et al. 2014). Consequently, an understanding of the space use and movement ecology of apex predators is key to effective conservation decision making for these species.

The jaguar (*Panthera onca*) is the largest feline in the Americas, distributed from the southwestern United States to northern Argentina, although it presently occupies <50% of its original range, and <80% of the range outside of Amazonia, due to habitat loss and persecution (Sanderson et al. 2002; Zellar 2007; de la Torre et al. 2017). Given the contraction of the species’ distribution, range-wide conservation efforts have focused upon maintaining connectivity among key populations throughout the species range (Sanderson et al. 2002; Rabinowitz and Zeller 2010), however, an effective implementation of this management approach is partly dependent upon a thorough understanding of the spatial and movement ecology of jaguars.

For a big cat the jaguar is relatively understudied (Brodie 2009), and although multiple studies have estimated jaguar home range size (Schaller and Crawshaw 1980; Rabinowitz and Nottingham 1986; Crawshaw and Quigley 1991; Crawshaw 1995; Scognamillo et al. 2002; Crawshaw et al. 2004; Silveira 2004; Cullen 2006; Azevedo and Murray 2007; Cavalcanti and Gese 2009; Tobler et al. 2013; Morato et al. 2016) and movements (Conde et al. 2010; Colchero et al. 2011; Sollman et al. 2011; Morato et al. 2016), there is still relatively little known about the species’ spatial and movement ecology. Since anthropogenic factors drive jaguar occurrence throughout its range by determining habitat availability and quality (Zeller et al. 2012; Petracca et al. 2014a,b; Thompson and Martinez 2015) this conspicuous knowledge gap on how jaguars perceive and use the landscape is of concern as it limits managers’ ability to quantifiably design and manage conservation landscapes for the jaguar.

Of further concern is that until recently jaguar home range estimates likely underestimated space use as VHF-based estimates were based upon small number of locations, while GPS-based estimates failed to account for autocorrelation inherent in GPS telemetry data (Morato et al. 2016). Furthermore, only recently have movement parameters and quantitative assessment of home range residency been estimated for jaguar (Morato et al. 2016).

Consequently, there is an important need for research that incorporates developing methodologies that account for and take advantage of autocorrelation in telemetry data to better quantify jaguar spatial and movement ecology.

Range-wide, the jaguar is considered near threatened (Caso et al. 2008), however, at the austral limit of its distribution the species is considered critically endangered in Argentina and endangered in Brazil and Paraguay. Although multiple studies have investigated space use by jaguar in Brazil and Argentina (Schaller and Crawshaw 1980; Crawshaw and Quigley 1991; Crawshaw 1995; Crawshaw et al. 2004; Silveira 2004; Cullen 2006; Azevedo and Murray 2007; Cavalcanti and Gese 2009; Morato et al. 2016) there has been no such research on the species in Paraguay despite a recognized need in the face of a rapid constriction in the species’ distribution in relation to a country-wide expansion of the agricultural sector (Secretaría del Ambiente et al. 2016) which has resulted in some of the highest rates of deforestation in the world (Hansen et al. 2013).

Given the status of the jaguar in Paraguay, the lack of information on the spatial and movement ecology of the species is of concern within the context of continued habitat loss, the maintenance of in-country and trans-boundary connectivity of populations, and their implications for the range-wide conservation of the jaguar. Consequently, we used GPS-based telemetry to study space use and movements of jaguars in western Paraguay in the Dry Chaco, Humid Chaco and Pantanal, the region with the largest jaguar population in the country. Moreover, we employed developing methodologies which allowed us to determine home range residency and account for autocorrelation in the data (Fleming et al. 2014, 2015; Calabrese et al. 2016), which in turn allowed for rigorous comparisons with estimates from other research employing the same methodologies (Morato et al. 2016).

Based upon carnivore ecology in general, and jaguar ecology specifically, we expected male home range and movement rates to be higher than females (Mikael 1989; Cavalcanti and Geese 2009; Conde et al. 2010; Sollmann et al. 2011; Morato et al. 2016) and that jaguars in the Dry Chaco would exhibit larger home ranges, higher movement rates, and more directional movement compared to those in the more productive habitats of the Humid Chaco and Pantanal (Mikael 1989; Fahrig 2007; Gutierrez-Gonzalez et al. 2012). Also, when comparing to other sites (Morato et al. 2016) we expected estimated from the Humid Chaco and Pantanal to be similar to those from the Brazilian Pantanal, while estimates from the Dry Chaco would be larger than those from more humid systems but possibly similar to jaguars from the Brazilian Cerrado due to biotic and abiotic similarities between systems. Apart from constituting an important contribution towards the conservation of jaguars within Paraguay, placing our results into a comparative context with research from neighboring countries will facilitate the efficacy of trans-boundary conservation efforts, with important implications for range-wide conservation strategies for jaguar.

## Materials and methods

### Study area

We conducted our study in three ecosystems in western Paraguay; Dry Chaco, Humid Chaco and Pantanal, (Figure 1). The Dry Chaco is comprised of xeric forest, savannas, and grasslands and the Humid Chaco and Pantanal are a mosaic of seasonally flooded grasslands, palm savanna and xerophilic woodlands on higher ground (Olson et al. 2001; Mereles et al. 2013). We note that delineations between the Humid Chaco and Pantanal differ (Olson et al. 2001; Mereles et al. 2013), however, for our purposes the similarities between systems and among our study sites in those systems make this discrepancy moot and consequently we treat the Humid Chaco and Pantanal as a single system in our analysis.

**Figure 1.**
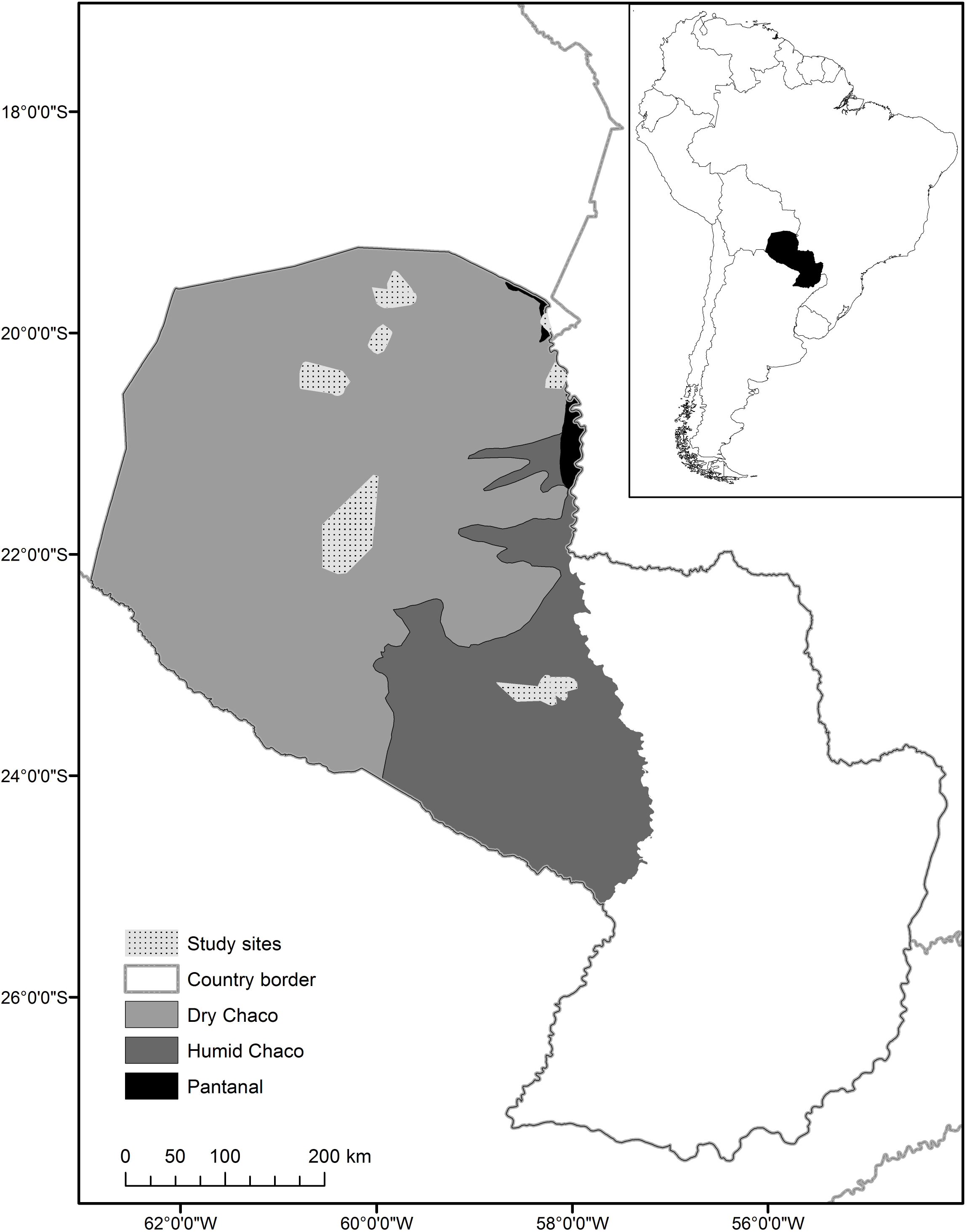
Map showing the distribution of the Dry and Humid Chaco and Pantanal in western Paraguay (Olson et al. 2001) and study areas where jaguar movements were monitored.

The western half of Paraguay is generally semi-arid with a pronounced east–west gradient in precipitation and humidity which divides the Chaco into the Humid Chaco with precipitation approximately > 1000 mm\year and the Dry Chaco with precipitation < 1000 mm\year (Olson et al. 2001). The Pantanal is also subjected to this east-west precipitation gradient; however, it and the Humid Chaco are also strongly effect by the hydrological cycles of the Rio Paraguay (Mereles et al. 2013).

In the Humid Chaco our study area was Estancia Aurora, a 30,000 ha cattle ranch in the north of the department of Villa Hayes and in the Pantanal on the 65,000 ha ranch Estancia Fortín Patria and on the 80,000 ha ranch Estancia Leda. In the central Dry Chaco, we worked on the 40,000 ha Faro Moro ranch and more northerly in the 7,200 km^2^ Defensores del Chaco National Park and the neighboring 269,000 ha of ranchland of the consortium *Grupo Chovoreca*.

### Jaguar captures

Jaguars were captured using trained hounds to tree or bay jaguars which were then anesthetized using a weight-dependent dose of a mix of ketamine hydrochloride and xylazine hydrochloride injected by a dart shot from a tranquilizer gun (McBride and McBride 2007). Capture methods followed ASM protocols (Sikes 2016) and in > 60 captures and recaptures of jaguar and puma over the study period there were no deaths or noticeable injury to animals.

From 2002-2009 jaguars were fitted with Telonics Generation II, data stored-on-board, GPS collars (Telonics, Mesa, Arizona, USA) which were set to record locations at 4 hour intervals. Starting in 2009 we used Northstar GPS collars (D-cell, Northstar, King George, Virginia, USA) programmed to record locations at three or four hour intervals and in 2012 we switched to Telonics Generation III GPS collars (Telonics, Mesa, Arizona, USA) which were set to record locations daily every two hours from 1800 to 0600 hours.

### Home range estimation

Semi-variogram analysis, model selection and AKDE estimates were undertaken using the *ctmm* package (Calabrese et al. 2016) in R 3.3.2 (R Development Core Team 2010). Starting values derived from semi-variograms were used for maximum likelihood model fitting with model selection based upon Akaike Information Criteria, adjusted for small sample size (AICc), and model weights (Fleming et al. 2014, 2015; Calabrese et al. 2016). We accounted for data collected with an irregular sampling schedule from collars used starting in 2012 with the *dt* argument within the *variogram* function in the *ctmm* package (Calabrese et al. 2016).

Movement models tested were an independent identically distributed (IID) model which ignores autocorrelation in the data and is equivalent to kernel density estimation (KDE) (Worton et al. 1989), a random search model (Brownian motion) with no home range, Brownian motion within a home range (Ornstein–Uhlenbeck, OU), and Ornstein–Uhlenbeck motion with foraging (OUF) (Fleming et al. 2014; Calabrese et al. 2016). Both the OU and OUF models produce estimates of home range size and home range crossing time, while the OUF model additionally estimates the velocity autocorrelation time scale (a measure of path sinuosity) and mean distance traveled (Fleming et al. 2014; Calabrese et al. 2016).

Home ranges were estimated using the best fit model for each individual using AKDE (Fleming et al. 2015; Calabrese et al. 2016). For comparison with home range estimates from previous research we estimated 95% KDE home ranges using the IID model and 95% Minimum Convex Polygons (MCP) home ranges using the adehabitatHR package in R (Calenge 2006) (Supplementary material Appendix 1).

### Core area estimation

We estimated core areas of AKDE home ranges as the area encompassed within the isopleth where the proportional use of the estimated home range is equal to the predicted probability of use (Seaman and Powell 1990; Bingham and Noon 1997; Vander Wal and Rodgers 2012). We determined this by fitting an exponential curve to the isopleths of the utility distribution of each individual at 10% increments from 10% to 90%, and at the 95% and 99% isopleths of the AKDE home range and the proportional area of the home range that each of those isopleths encompassed based upon the area of the 99% home range estimate. We then determined the threshold where proportional home range size begins to increase at a rate greater than the probability of use (slope=1; Seaman and Powell 1990; Bingham and Noon 1997; Vander Wal and Rodgers 2012) to define the isopleth that represented the core area boundary.

### Statistical analyses

For our statistical analysis we combined jaguars from the Humid Chaco and the Pantanal into a single group as the characteristics of the system are highly similar, the delineation between the two systems is debatable (Olson et al. 2001; Mereles et al. 2013), and consequently jaguars from those systems are subjected to similar ecological and anthropogenic drivers. Additionally, only individuals that exhibited residency in their movement behavior through semi-variogram analysis and space use best explained by the OUF model were included in our comparative analysis of differences between sexes and ecosystems.

We used a fixed-effect one-way analysis of variance (ANOVA) in a Bayesian modeling framework to test for differences in estimates of home range size, home range crossing time, directionality in movement (velocity autocorrelation time scale) and mean daily distance traveled between sexes across systems, between systems (sexes combined), between sexes within a system, and between same sexes between systems. We tested normality using the Shapiro-Wilk test and log-transforming the data when its distribution did not meet assumptions of normality.

All analyses were undertaken in R 3.2.2. (R Development Core Team 2010) using WinBUGS (Lunn et al. 2000) and the *R2bugs* package (Sturtz et al. 2005) for the Bayesian analysis. We ran 3 chains in WinBUGS with 100,000 iterations and a 20,000 iteration burn-in period; confirming convergence by a scale reduction factor ≤1.01 and visual inspection of trace plots for lack of autocorrelation. We tested differences between groups by taking 10,000 random samples from posterior distributions for each group of interest, comparing the proportional frequency that posterior estimates parameters were greater for males than females overall and within systems, greater for all individuals, and between same sexes, in the Dry Chaco compared to the Humid Chaco/Pantanal.

## Results

### Jaguar captures and data collection

We captured and collared 35 jaguars from June 2002 to June 2014 of which 19 individuals provided sufficient data for analysis; 7 in the Dry Chaco (5 males, 2 females), 9 in the Humid Chaco (3 males, 6 females) and 3 in the Pantanal (1 male, 2 females) with estimated ages between 2 and 10 years (Table 1). Collars collected data between 52 and 439 days, obtaining from 148 to 3462 locations (Table 1). The length of the study period and the annual frequency of captures were dependent upon resource availability and logistical restraints that dictated captures and collar recovery.

**Table 1.** Sex, age, sample characteristics and estimated movement parameters, AKDE home range, core area and core area utility distribution isopleths for study jaguars in the Paraguayan Dry Chaco, Humid Chaco and Pantanal.

| ID | Sex/age<br>(yr) | Number<br>of<br>fixes/days | Velocity<br>autocorre<br>lation<br>timescale<br>(h) | Home<br>range<br>crossing<br>time<br>(days) | Average<br>distance<br>traveled<br>(km/day) | Home range (km<sup>2</sup>)<br>(95% CI) | Core<br>(km<sup>2</sup>) | Core area<br>isopleths<br>(%) |
| --- | --- | --- | --- | --- | --- | --- | --- | --- |
| *Dry Chaco* |  |  |  |  |  |  |  |  |
| DC1 | M/5 | 1094/376 | 1.1 | 8.0 | 28.8 | 2143 (1558-2820) | 504 | 59 |
| DC3 | M/2 | 722/363 | 1.8 | 11.5 | 7.9 | 421 (288-580) | 107 | 63 |
| DC4 | M/5 | 620/82 | 1.9 | 3.5 | 15.0 | 550 (349-797) | 182 | 58 |
| DC6 | M/7 | 1387/393 | 2.2 | 4.8 | 19.3 | 1063 (822-1335) | 329 | 57 |
| DC7 | M/5 | 3462/439 | 1.4 | 2.7 | 17.1 | 445 (381-515) | 85 | 64 |
| DC5 | F/6 | 1610/386 | 1.1 | 11.5 | 11.8 | 591 (411-805) | 178 | 59 |
| DC2 | F/8 | 921/379 | 1.7 | 9.5 | 9.7 | 511 (363-683) | 176 | 56 |
| *Humid Chaco/Pantanal* |  |  |  |  |  |  |  |  |
| Pan2 | F/2 | 1694/375 | 1.1 | 4.3 | 7.9 | 71 (58-85) | 24 | 60 |
| HC5 | F/4 | 593/242 | 0.5 | 1.2 | 20.9 | 92 (75-110) | 23 | 61 |
| HC4 | F/3 | 288/266 | 1.5 | 6.2 | 9.3 | 270 (187-369) | 86 | 57 |
| HC8 | F/1 | 980/170 | 0.2 | 10.2 | 13.7 | 121 (71-183) | 32 | 58 |
| HC7 | F/6 | 1668/324 | 0.1 | 9.2 | 22.3 | 246 (172-332) | 73 | 57 |
| HC9 | F/6 | 928/362 | 0.2 | 9.7 | 13.9 | 118 (83-159) | 33 | 59 |
| Pan3 | M/6 | 727/192 | 1.4 | 3.4 | 16.6 | 428 (320-550) | 134 | 57 |
| HC3 | M/4 | 983/143 | 1.4 | 4.4 | 15.0 | 424 (290-584) | 138 | 56 |
| HC6 | M/10 | 660/133 | 0.9 | 5.5 | 13.4 | 341 (216-494) | 91 | 60 |
| Pan1 | F/4 | 1695/366 | NA | 3.5 | NA | 550 (349-797) | 21 | 60 |
| HC1 | M/6 | 148/88 | NA | 5.9 | NA | 958 (534-1505) | 283 | 58 |
| HC2 | F/6 | 280/54 | NA | 5.7 | NA | 73 (35-125) | 22 | 57 |

### Home range, core area and movement parameter estimates

Best fitting models for the movement of jaguars were either the OU or OUF models with 16 individuals demonstrating residency (Table 1). Estimated home range sizes varied between 86 and 2,909 km^2^ and core areas between 21-509 km^2^. Core areas were represented by a consistent proportion of the utility distribution; ranging between 56%-64% isopleths (Table 2).

**Table 2.** Probabilites, based upon posterior distributions, that home range and movement parameters are different between sex and ecosystem, between sexes within systems, and between same sexes between systems.

|  | Home range<br>(km<sup>2</sup>) | Home range<br>crossing time<br>(days) | Velocity<br>autocorrelation<br>timescale (h) | Average distance<br>traveled<br>(km/day) |
| --- | --- | --- | --- | --- |
| Dry Chaco male > Dry<br>Chaco female | 0.71 | 0.08 | 0.71 | 0.89 |
| Humid<br>Chaco/Pantanal male ><br>Humid<br>Chaco/Pantanal female | 0.99 | 0.17 | 0.95 | 0.52 |
| Dry Chaco male ><br>Humid<br>Chaco/Pantanal male | 0.91 | 0.75 | 0.86 | 0.72 |
| Dry Chaco female ><br>Humid<br>Chaco/Pantanal female | 0.99 | 0.89 | 0.96 | 0.23 |
| All Dry Chaco > All<br>Humid<br>Chaco/Pantanal | 1 | 0.77 | 0.99 | 0.61 |
| Male > Female | 0.99 | 0.10 | 0.99 | 0.84 |

Male and female mean home range size were 727 km^2^ (95% CI:355-1954) and 255 km^2^ (95% CI:90-578), respectively and 818 km^2^ (95% CI:425-1981) and 237 km^2^ (95% CI:90-427) for jaguars in the Dry Chaco and Humid Chaco/Pantanal, respectively (Fig. 2, Fig.3). In the Dry Chaco mean home range size for males was 925 km^2^ (95% CI:424-2035) and 551 km^2^ (95% CI:513-590) for females, while in the Humid Chaco/Pantanal the mean home range was 398 km^2^ (95% CI:345-427) and 156 km^2^ (95% CI:90-267) for males and females, respectively (Fig. 4).

**Figure 2.**
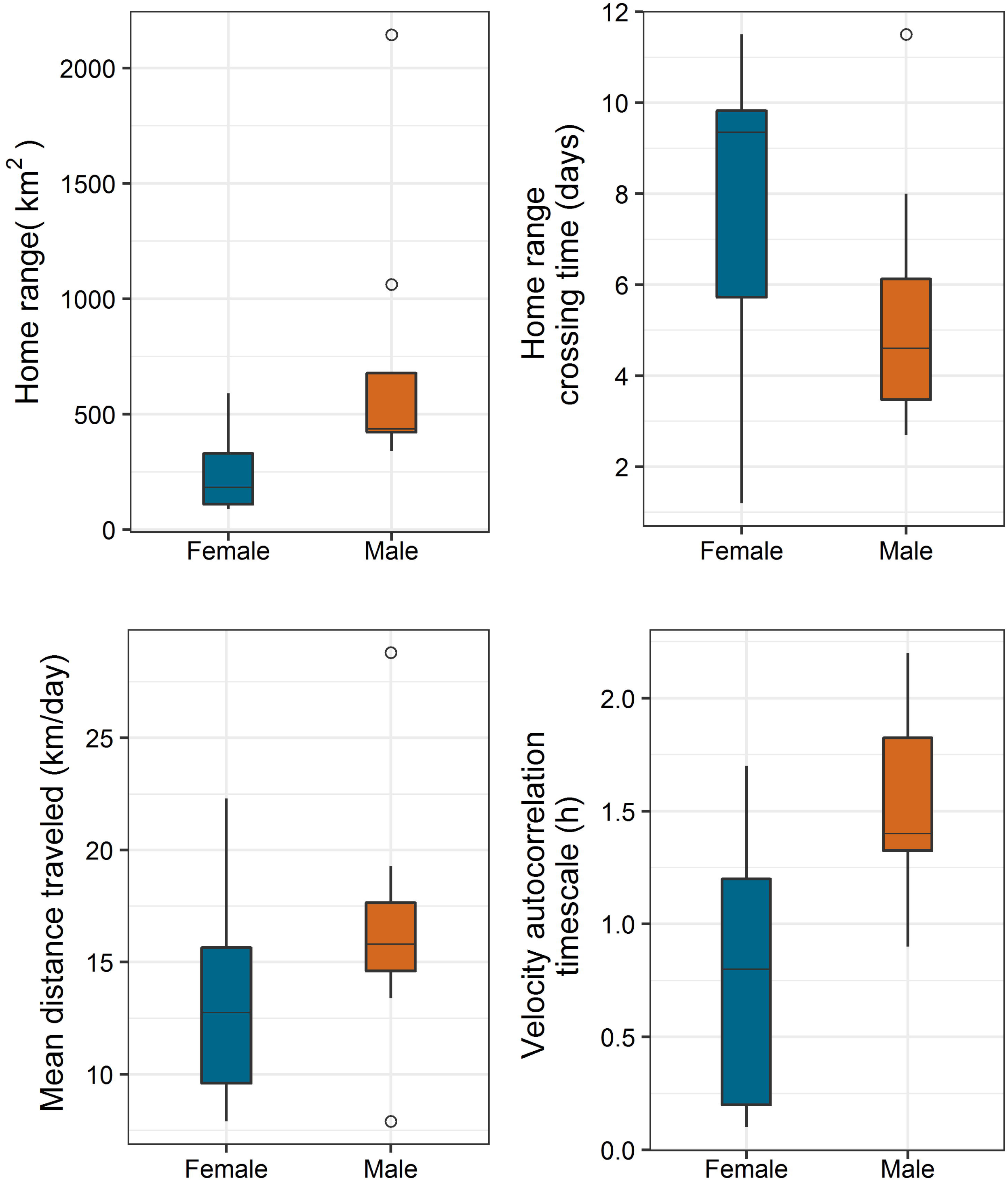
Home range and movement parameters of male and female jaguars across all study sites.

**Figure 3.**
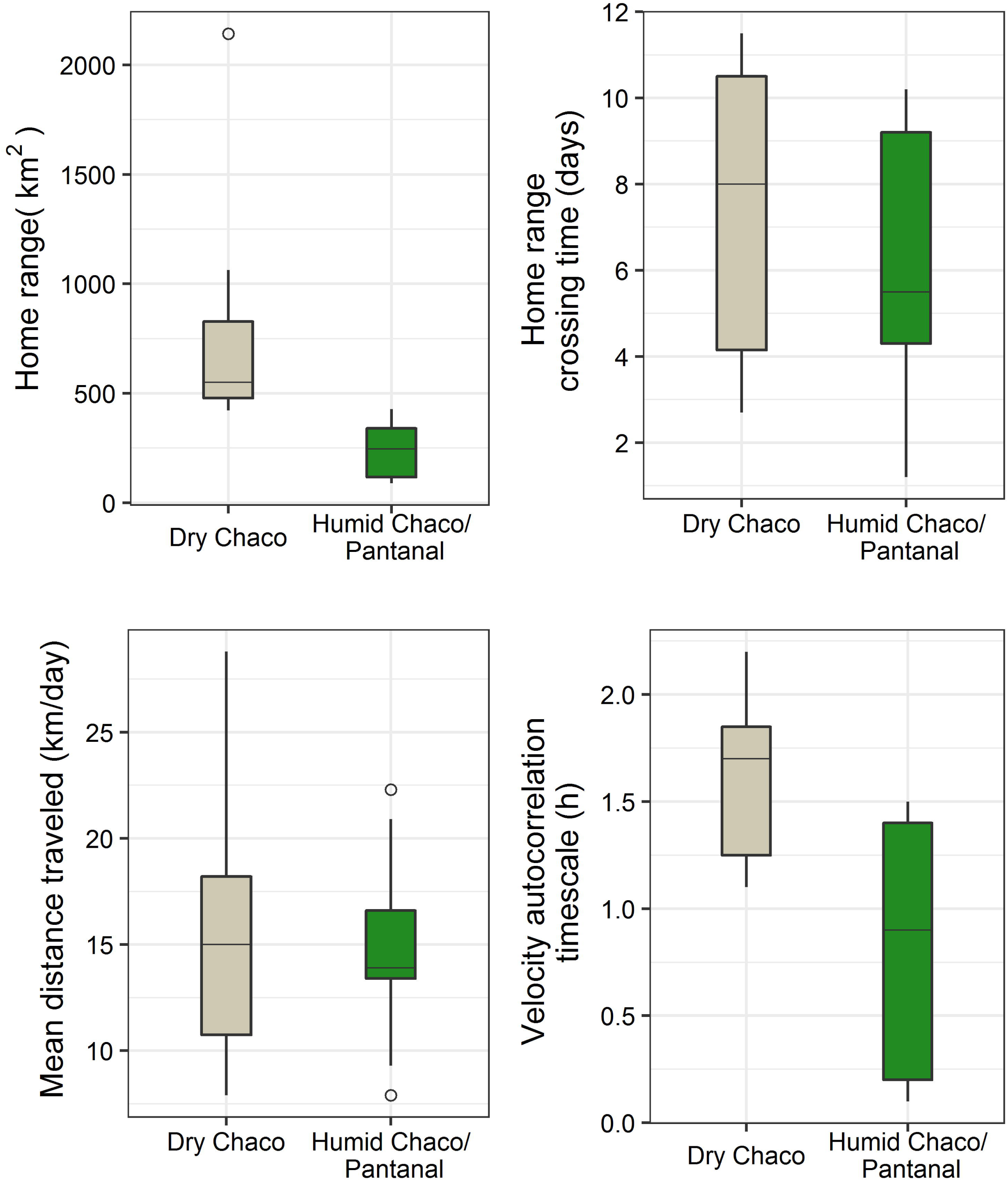
Home range and movement parameters for study jaguars in the Dry Chaco and Humid Chaco/Pantanal.

**Figure 4.**
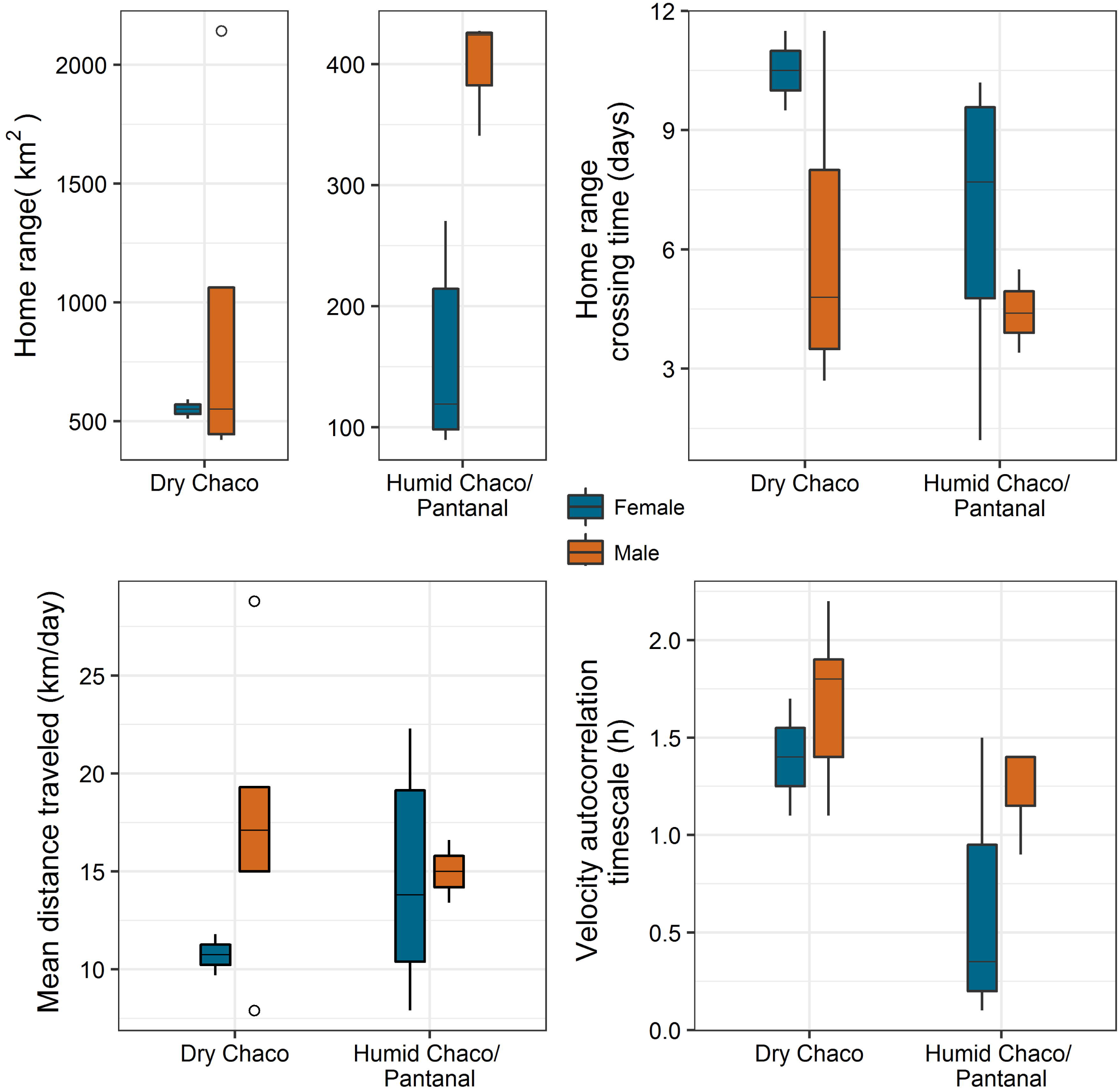
Home range and movement parameters of male and female jaguars in the Dry Chaco and Humid Chaco/Pantanal.

Males demonstrated larger home ranges (*P*=0.99), higher daily movement (*P*=0.84), greater directionality in movement (velocity autocorrelation time scale) (*P*=0.84) and lower home range crossing times (*P*= 0.9) (Table 2, Figure 2). Between systems, home ranges were larger (*P*=1), movements more directional (*P*=0.99) and home range crossing times greater (*P*=0.77) in the Dry Chaco, while daily travel distance was similar between systems but with a slightly higher probability of being larger in the Dry Chaco (*P*= 0.61, Figure 3).

Between systems males in the Dry Chaco had higher probabilities to have larger home ranges (*P*=0.91), higher home range crossing time (*P*=0.75), greater directionality in movement (*P*=0.86), and greater daily travel distances (*P*=0.72) (Table 2), although values for all parameters were more variable in males from the Dry Chaco (Figure 4). A similar pattern was evident between females in both systems for home range size (*P*=0.99), home range crossing time (*P*=0.89) and directionality in movement (*P*=0.96) which were greater for females in the Dry Chaco, however, females in the Dry Chaco had lower daily movements (*P*=0.23) than those in the Humid Chaco/Pantanal (Table 4).

## Discussion

We present the first estimates of movement parameters and home range and core area for jaguar in the Dry Chaco, Humid Chaco, and Paraguayan Pantanal, which furthermore take advantage of developing methods to empirically test for home range residency and account for autocorrelation in telemetry data when estimating space use (Fleming et al. 2014. 2015; Calabrese et al. 2016). Our results include the largest home range estimates recorded for jaguar (Dry Chaco) and, as expected, jaguars in the more productive Humid Chaco/Pantanal had smaller home ranges, lower movement rates and had less directionality in movements compared to jaguars in the Dry Chaco. Also, consistent with previous research males had larger home ranges, higher movement rates and more directional movements than females overall and within systems.

Overall and between systems male home ranges were larger than females which was expected (Calvalcanti and Gese 2009; Sollmann et al. 2011; Morato et al. 2016) as smaller home ranges of females are driven by food availability in relation to reproductive and offspring rearing needs which in-turn drives larger male home ranges towards optimizing reproductive opportunities (Mikael 1989; Sunquist and Sunquist 1989). This relationship is further supported by our estimated movement parameters which showed that males traveled farther, faster, and more directionally than females in utilizing home ranges.

Consistent with our expectations home range sizes of jaguars in the Dry Chaco were larger than in the Humid Chaco and Pantanal, overall and between sexes within systems where male home ranges were greater than females. The larger home ranges in the Dry Chaco are attributable to the lower productivity of that semi-arid ecosystem, more heterogeneously distributed prey and water, and negative effects of anthropogenic factors (i.e., deforestation; Fahrig 2007; Gutierrez-Gonzalez et al. 2012). The difference between sexes within systems is attributable to differences in territorial organization stemming from aforementioned reproductive and social needs (Mikael 1989; Sunquist and Sunquist 1989).

Home range estimates from the Dry Chaco for both males and females are considerably larger than other estimates from this study and Morato et al. (2016), although our estimates of male home range size from the Dry Chaco (mean:925 km^2^, 95% CI:424-2035) are consistent with the estimate for a single male from the Brazilian Cerrado (1269 km^2^), a semi-arid ecosystem with environmental and land use similarities to the Gran Chaco. Morato et al. (2016) demonstrated that increasing home range size of jaguars was associated with lower habitat quality, which is consistent with the very large home ranges from the Dry Chaco which were closest in size to Morato et al’s (2016) home ranges in the Atlantic forest which they considered to be of the lowest habitat quality of their study areas.

We expected home range sizes from the Humid Chaco/Pantanal to be similar to estimates from the Brazilian Pantanal, however, our estimates were 59% and 112% larger for males and females, respectively than home ranges reported for the Brazilian Pantanal; falling between estimates from the Amazon and Atlantic forest, although most similar to jaguars from the Amazon (Morato et al. 2016; Fig. 5). These differences may be related to differences in the geomorphology of the two regions and its interaction with the local hydrological cycles.

**Figure 5.**
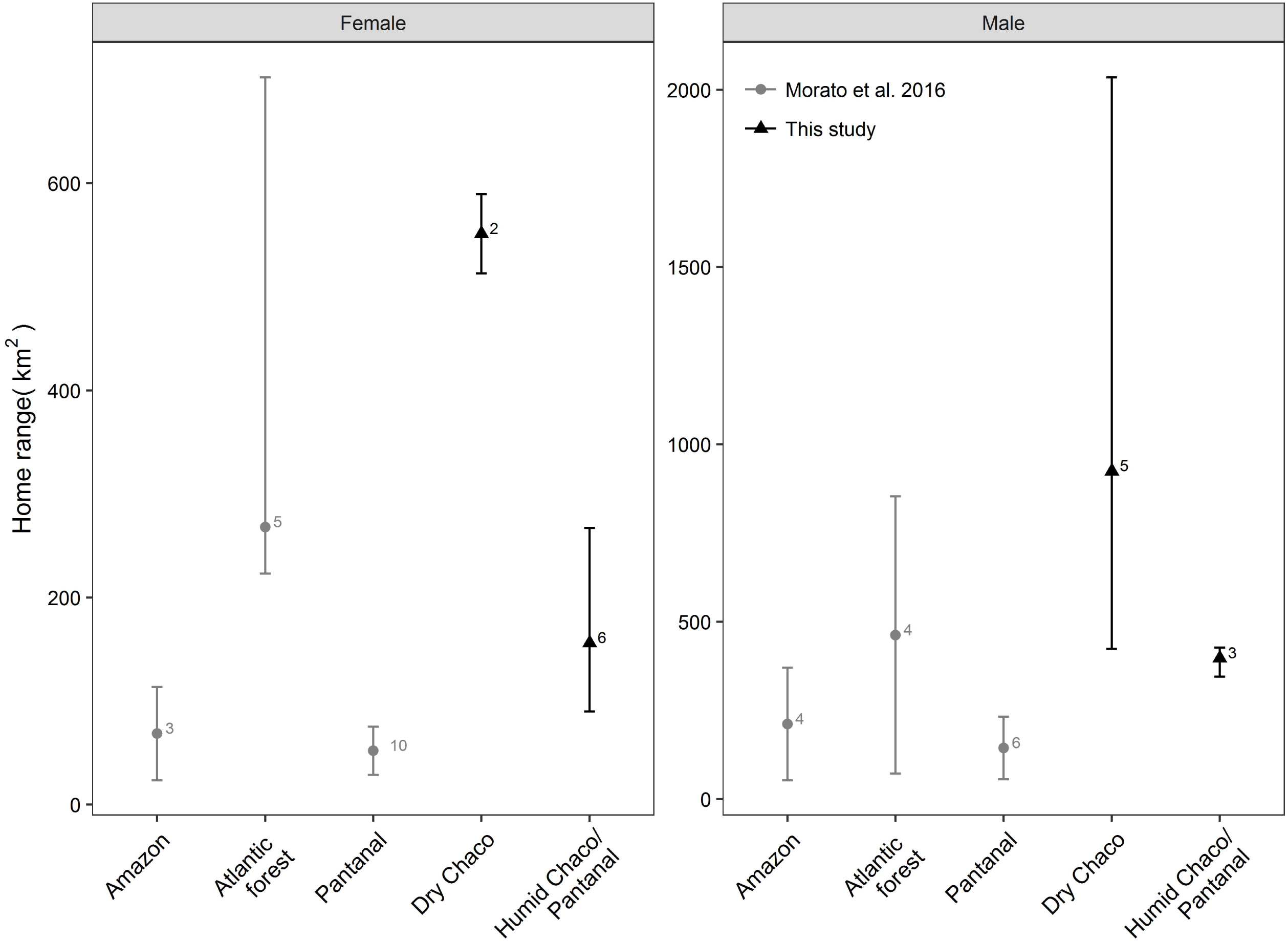
Mean male and female home ranges (error bars represent 95% confidence interval) from this study and AKDE estimates from Morato et al. (2016) by ecosystem. The numbers next to points represent sample size.

The Paraguayan Pantanal and our study area in the Humid Chaco have less forest area and a relatively greater area of inundated land during a large portion of the year compared to the Pantanal study areas of Morato et al. (2016) in Brazil. Consequently, the reduced forest area, with smaller and more isolated forest patches during annual flooding, could drive the comparatively larger home ranges observed in the Paraguayan Pantanal and Humid Chaco, although reduced jaguar densities resulting from persecution may also play a role in liberating available space and permitting greater space use.

Differences in the mean movement parameters were evident between jaguars in the Humid Chaco/Pantanal and in the Brazilian Pantanal whereby movements were more directional in the Humid Chaco/Pantanal, although still relatively sinuous but most similar to jaguars in the Atlantic forest, while daily movements were very similar to those in the Amazon (Fig. 6). Jaguars in the Dry Chaco had high movement rates and directionality in movement, similar to individuals in the Amazon from seasonally flooded forests (Morato et al. 2016).

**Figure 6.**
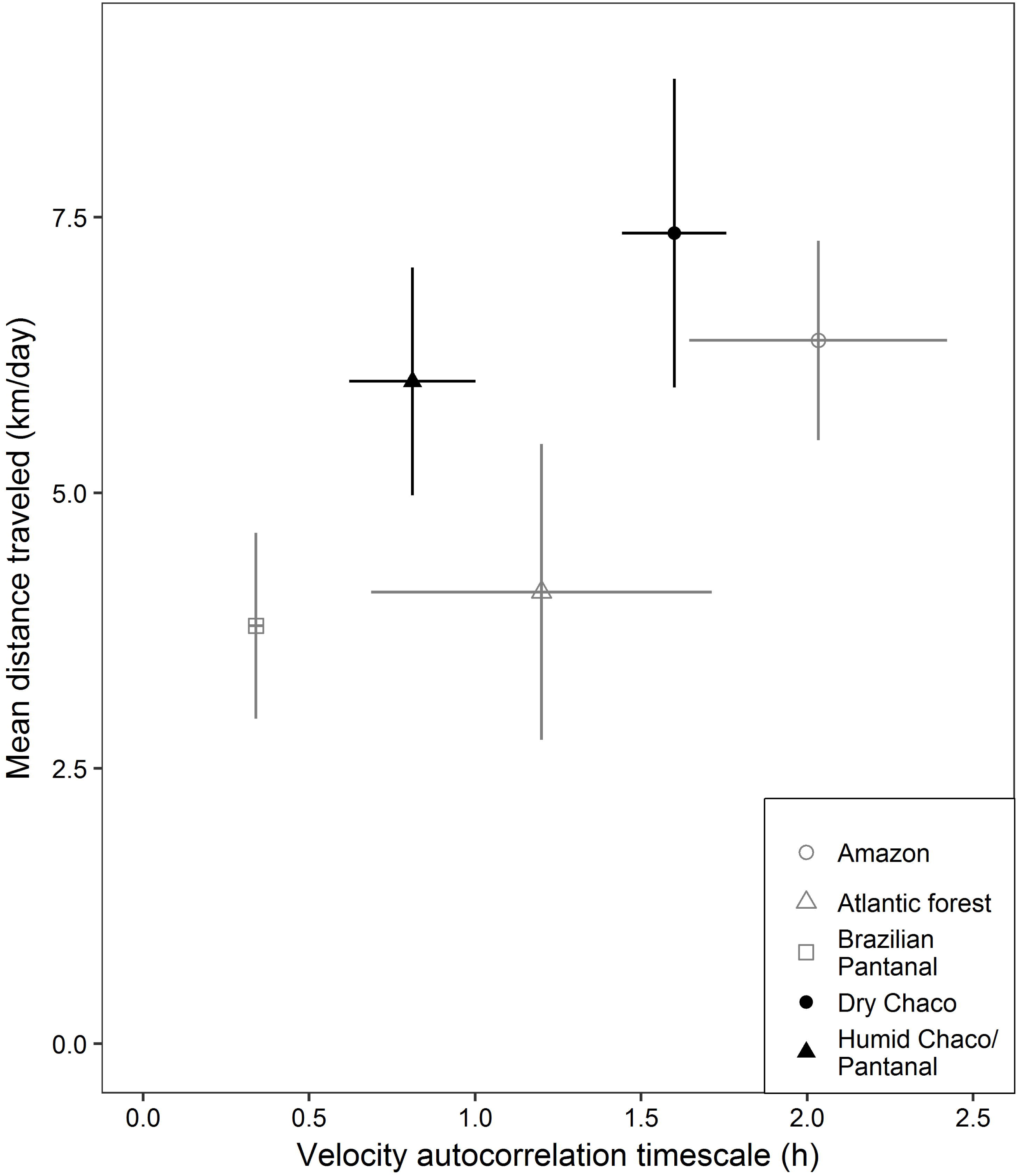
Mean of the mean distance traveled and the velocity autocorrelation timescale (error bars represent SE) of jaguars from this study and mean estimates from Morato et al. (2016) by ecosystem.

We believe that these similarities are responses to movements among sporadically distributed critical resources despite the large differences in ecosystem characteristics.

Conversely, although daily movement rate of jaguars in the Humid Chaco/Pantanal were similar to those in the Dry Chaco and Amazon, the relatively low directionality demonstrated by jaguars in the Humid Chaco/Pantanal suggests that, although jaguars are covering relatively large areas, movements are in response to more homogenously distributed resources within home ranges.

Core areas, as measured by the utility distribution isopleth were highly similar across systems and sexes, encompassed on average by the 59% isopleth (95% CI:56-64%), which represented on average 29% (95% CI:21-34%) of total home range area. This indicates that despite home range size, sex, or system jaguars are most intensively using about a third of their home range area. Additionally, our results suggests a cautious interpretation of arbitrarily defined core area delimitations, typically assigned to the 50% utility distribution isopleths which falls outside of the 95% confidence limits of our estimates (Powell 2012).

In light of the extensive deforestation that is occurring in the Dry Chaco of western Paraguay, the large home ranges that we observed in this system, which are consistent with the estimated low density of jaguar in the Bolivian Dry Chaco (Noss et al. 2012), are of concern as they demonstrate the large forested area that jaguars in the Dry Chaco require. In the Humid Chaco/Pantanal spatial requirement of jaguars were greater than expected based on estimates from the Brazilian Pantanal, which suggests lower than expected densities in these systems in Paraguay and cautions against extrapolating population parameter estimates from other regions within the Pantanal to the Rio Paraguay flood plain in Paraguay.

In both the Dry Chaco and the Humid Chaco/Pantanal we recognize that there may be an important effect on space use caused by reduced jaguar densities from persecution which is pervasive throughout western Paraguay, illustrated by our confirmation, or high probability, of ∼75% of our study animals being killed due to persecution. Persecution is common throughout the range of the jaguar, however, its practice and magnitude is not equivocal geographically and consequently how the removal of individuals may impact space use, and subsequently comparisons among ecosystems and regions, needs to be considered and is of interest for future research.

The large spatial requirements of jaguars in western Paraguay, particularly in the Dry Chaco, indicate that the protected areas of the region which, represent <5% of the total regional area are likely insufficient to maintain a viable regional population, especially in light of the level of persecution on private lands. This highlights an urgent need to mitigate jaguar-human conflict in the region by actively including the livestock production sector in the conservation decision making process. Furthermore, given continuing deforestation, conservation initiatives need to take into account the large spatial needs of jaguar in western Paraguay by recognizing and incorporating the role of private lands in the long-term conservation of the species in Paraguay and in maintaining trans-boundary connectivity among populations.

## Acknowledgements

We thank DVM Sybil Zavala, Cougar McBride and Caleb McBride for assistance in the field and the many sportsman and conservationist who contributed to supporting this work. This research was conducted under the permission of the Secretariat of the Environment (SEAM) of Paraguay. JJT was supported by the Consejo Nacional de Ciencia y Tecnología of Paraguay (CONACYT).

## Supplementary material

**Table A.1.**
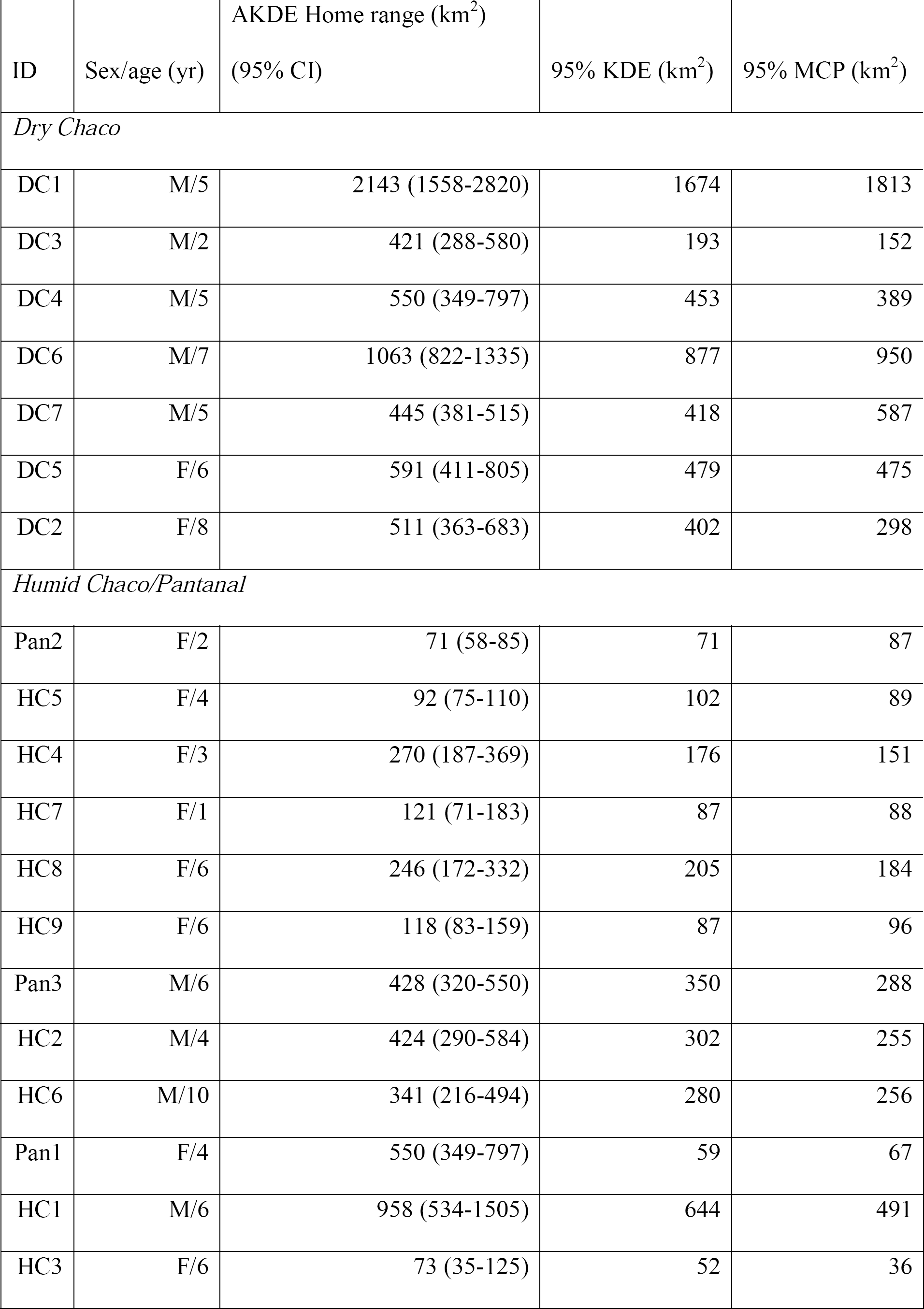
Comparative home range sizes for study jaguars based upon autocorrelated kernel density estimator (AKDE), 95% kernel density estimator (KDE), and 95% minimum convex polygon (MCP).

| ID | Sex/age (yr) | AKDE Home range (km<sup>2</sup>)<br>(95% CI) | 95% KDE (km<sup>2</sup>) | 95% MCP (km<sup>2</sup>) |
| --- | --- | --- | --- | --- |
| *Dry Chaco* |  |  |  |  |
| DC1 | M/5 | 2143 (1558-2820) | 1674 | 1813 |
| DC3 | M/2 | 421 (288-580) | 193 | 152 |
| DC4 | M/5 | 550 (349-797) | 453 | 389 |
| DC6 | M/7 | 1063 (822-1335) | 877 | 950 |
| DC7 | M/5 | 445 (381-515) | 418 | 587 |
| DC5 | F/6 | 591 (411-805) | 479 | 475 |
| DC2 | F/8 | 511 (363-683) | 402 | 298 |
| *Humid Chaco/Pantanal* |  |  |  |  |
| Pan2 | F/2 | 71 (58-85) | 71 | 87 |
| HC5 | F/4 | 92 (75-110) | 102 | 89 |
| HC4 | F/3 | 270 (187-369) | 176 | 151 |
| HC7 | F/1 | 121 (71-183) | 87 | 88 |
| HC8 | F/6 | 246 (172-332) | 205 | 184 |
| HC9 | F/6 | 118 (83-159) | 87 | 96 |
| Pan3 | M/6 | 428 (320-550) | 350 | 288 |
| HC2 | M/4 | 424 (290-584) | 302 | 255 |
| HC6 | M/10 | 341 (216-494) | 280 | 256 |
| Pan1 | F/4 | 550 (349-797) | 59 | 67 |
| HC1 | M/6 | 958 (534-1505) | 644 | 491 |
| HC3 | F/6 | 73 (35-125) | 52 | 36 |

